# Identification and Expression of the peach TNL *RMia* genes for the Resistance to the Root-knot Nematode *Meloidogyne incognita*

**DOI:** 10.1101/2024.10.16.618674

**Authors:** Henri Duval, Laure Heurtevin, Naïma Dlalah, Claire Caravel, Caroline Callot, Cyril Van Ghelder

## Abstract

Root-Knot Nematodes (RKN) are major pests for world agriculture. As RKN are polyphagous endoparasites, present in various climates and able to multiply rapidly, the quest to discover new natural resistances in plants is essential. In *Prunus* species, several major resistance genes to RKN that belong to the NLR family, have been detected in recent years. In Peach, the RMia gene, which confers resistance to the RKN *Meloidogyne incognita, M. arenaria*, and *M. ethiopica*, was localized on the chromosome 2. In this study, we used short-and long-read sequencing technologies to identify the RMia gene in two independent RKN-resistant peach varieties. We characterized three candidate genes and studied their expression profile upon RKN infection. Our study identified and characterized these candidates that are most likely involved in the RKN resistance. Ultimately, we developed new molecular markers associated to this resistance that can be used in breeding programs.

## 1. Introduction

Among the plant parasitic nematodes, Root-Knot Nematodes (RKN), *Meloidogyne spp*., are often described as the most important genus in terms of scientific studies and caused damages (Jones et al. 2013). The model species *M. incognita* together with *M. javanica, M. arenaria, M. hapla* and the emerging species, *M. floridensis* and *M. enterolobii* represent a threat for agriculture due to their host range, distribution, and life traits. Indeed, these telluric pathogens have a worldwide distribution, are able to infect almost all the cultivated plant species, and can complete their life cycle in a few weeks producing a thousand juveniles per female stage in the model species *M. incognita* for instance (Abad et al. 2008). With the progressive removal of the most harmful nematicides, other alternatives to control these pests are required. The combination of innovative cultural methods, the use of biocontrol agents and the reasoned utilization of resistant plant cultivars is the most promising approach to control RKN and minimizing harmful effects on the environment (Djian-Caporalino et al. 2019). Therefore, the identification and characterization of major Resistance genes (R genes) against RKN are crucial in this context. As the risk of R gene circumvention increases in perennial plants, due to longer contact between host and pathogens, the opportunity to combine several R genes in one cultivar (*i.e*. gene pyramiding) reinforces resistance’s durability. This approach is particularly relevant for the *Prunus* genus that gathers many species of agronomical interest, including peach, plum and almond, which are susceptible to RKN. As hybrids and grafts can be carried out between *Prunus* species, investigating the *Prunus* R genes repertoire is particularly promising.

NLR genes or Nucleotide Binding and Leucine Rich-Repeat receptor genes (NB-LRR) represent a major resistant gene family in plants. They are composed of 3 different classes mainly characterized by a conserved structure made of the Nucleotide Binding and the Leucine Rich-Repeat domains, and different N-terminal domains. The CNL (including a N-terminal Coiled-Coil domain), the RNL (RPW8 domain) and the TNL (TIR or Toll-like Interleukin Receptor-1 domain) form this gene superfamily. Although NLR numbers are numerous in plant genomes (*i.e*. often >100 per genomes), the ratio between the three classes greatly varies in plant genomes reflecting various defense strategies (Shao et al. 2014). NLR are intracellular receptors that can detect the presence of pathogens through the direct or indirect recognition of molecules (grouped under the name of effectors) secreted by invaders to feed on plants and escape plant immunity. This detection is often carried out by C-terminal domains of NLR sensors which will activate, through polymerization and N-terminal domains, other NLR helpers to trigger a downstream signaling pathway leading to cell death. Thus, NLR act in complex networks to confer plant immunity (Castel et al. 2019). Therefore, NLR are pivotal factors of the so-called Effector-Triggered Immunity (ETI) in plants. Recently, different TNL models have been structurally studied, revealing that TNL C-terminal domains (*i.e*. Leucine Rich Repeat and Post-LRR C-JID domains) directly bind the corresponding effector. Then, a superstructure (resistosome), made of a tetramerization of the TNL sensor, will lead to cell death through a mechanism that still needs to be more documented (Ma et al. 2020; Martin et al. 2020)

In the past two decades, several RKN resistance genes have been detected and identified in *Prunus* species (Saucet et al. 2016). In plum, the dominant TNL Ma gene, cloned in 2008, is a sole R gene that confers a full resistance to all *Meloidogyne* species tested including the emerging species *M. enterolobii* and *M. floridensis* (Duval et al. 2019). To date, this gene, displaying a unique structure among TNL, is not circumvented by high and durable inoculum pressure or by high temperature (Claverie et al. 2011). *Meloidogyne* juveniles will be able to penetrate the roots of the Ma-carrier accession, *P. cerasifera* P2175, however they will not induce any feeding site, preventing any nematode development. Following nematode recognition, the Ma gene will induce an Hypersensitive-like Reaction around RKN juveniles (Khallouk et al. 2011). Two genes, Rjap and RMja, which are most likely the orthologs of the Ma gene in *P. salicina* and in *P. dulcis*, respectively, display different spectra of resistance. While the Rjap gene has the same broad-spectrum of resistance as Ma, RMja does not confer a resistance to *M. incognita and M. floridensis*, making it an interesting model to study (Van Ghelder et al. 2018; Duval et al. 2019). Besides, two other TNL genes (PsoRPM2 and PsoRPM3) isolated from the wild myrobalan plum (*Prunus sogdiana*) conferred a resistance to *M. incognita* when transferred to tobacco (Zhu et al. 2017; Xiao et al. 2021).

In peach, the RMia gene represents a major source of resistance to RKN as it confers a resistance to *M. incognita, M. arenaria* and *M. ethiopica* (Duval et al. 2019). In a previous study, the gene has been localized in a region of approximately 92Kb flanked by 2 SNP markers (SNP_APP91 and SNP_APP92, named hereafter SP91 and SP92) using a high resolution mapping approach based on 790 hybrids involving the RMia carrier, *P. persica var. Nemared* (Duval et al. 2014). Interestingly, the unrelated variety *P. persica var. ‘Weeping Flower Peach’ (WFP)*, which contains the resistance gene (Rm1) to the aphid *Myzus persicae*, also contains the RMia gene (Duval et al. 2022). Considering that Ma and RMja genes are now identified orthologs localized in the Linkage Group 7 of the peach genome, the identification and the development of molecular markers for the RMia gene is therefore a crucial step for breeders to consider pyramiding of both sources of resistance. Besides, the comparison of sequences and resistance spectra between the RMia gene and the Ma and RMja genes, would be of great interest to shed light on key molecular determinants involved in RKN resistance in plants.

In this study, we obtained the genomic sequence lying between two SNP markers in two different peach varieties carrying the RMia gene. We performed an *in silico* structural and functional annotation of the region of interest supported by long-read transcriptomic data. We took this opportunity to develop reliable markers that may be of interest for breeding programs. Finally, we explored the expression pattern of the three candidate genes, which belong to the TNL family, upon RKN infection. We observed a pattern of expression that differs between TNL but conserved between both resistant peach varieties, thus revealing a possible intricate stress response. By revealing this cluster of resistance, our study paves the way for future functional studies in a species of high agronomic interest.

## 2. Materials and Methods

### 2.1 Plant Materials

The RKN-resistant peach clones, S2678 Weeping Flower Peach (WFP) and Nemared, were maintained in the INRAE collection in pots under insect-proof tunnels. The clone S2678 (WFP) was introduced in the INRAE collection from Clemson University. This clone has red double flowers, a weeping growth habit, and is resistant to the aphids *Myzus persicae* and *Myzus* variant (Massonie and Maison 1979; Duval et al. 2022). Nemared was released by USDA Fresno, California and originated from F3 seedlings of ‘Nemaguard’ x ‘a red-leaf seedling’ (Ramming and Tanner 1983). The plants used for the RKN infestation tests were produced by cuttings issued from WFP and Nemared. We also tested seeds of one hybrid ‘PRMN Z64P40’ obtained from a self-pollination of one nematode-resistant hybrid of the cross between of three susceptible peach variety and Nemared: [(Pamirskii x Rubira) x (Montclar x Nemared)]. We genotyped the peach varieties and hybrids using the markers developed in this study. These peach accessions have been phenotyped for their response to RKN either in previous studies or in this study (**Table S1**).

### 2.2 Root-knot nematodes

RKN, *M. incognita*, was multiplied using susceptible tomato plants, *Solanum lycopersicum* cv. ‘St Pierre’, in a greenhouse (temperature range 25-30°C). The nematode identity was verified by esterase profiling. After six weeks, tomato roots were collected and washed. The root system, full of galls, was put onto a sieve inside a bucket, and the whole system was placed in a mist chamber for 2-3 days. Freshly hatched juveniles stage 2 (J2) were collected in tap water and counted. Approximately 10 000 J2 were used to inoculate each plant of *P. persica* cv. ‘WFP’ and *P. persica* hybrid ‘PRMN Z64P40’. Both accessions carried the RMia gene. *M. incognita* J2 were distributed in four holes around the base of the plant to ensure homogeneous inoculation. The inoculation was carried out in a greenhouse under controlled conditions on young plants grown in small pots. Control plants, free of nematodes, were placed in the same growth conditions. After 10 days, roots of healthy control and nematode-infected plants were collected for subsequent RNA extraction.

### 2.3 Gene Expression (qPCR)

10 days after inoculation, root apexes from nematode-inoculated and control of WFP and ‘PRMN Z64P40’ plants were retrieved and flash frozen for storage. Total RNA was extracted from frozen samples using the NucleoSpin^®^RNA Plant Kit (Macherey-Nagel, Dueren, Germany) according to the manufacturer’s instructions. RNA yield and purity were assessed using a Nanodrop^®^ by spectrophotometric absorbance and agarose gel electrophoresis. In total, 500 μg of cDNAs were synthesized using the Affinity Script Multiple Temperature cDNA Synthesis Kit (Agilent), following the instructions provided. Expression analysis was performed by quantitative real-time PCR (qRT-PCR) using a Stratagene® Mx3005P® qPCR System instrument and Brilliant III SYBR Green qPCR Master Mix with Low ROX (Agilent Technologies). The reaction was prepared with each primer (0.5 μL, 10 μM), 2 μL of 1:10 diluted cDNA, RNase-free water (5 μL), and Brilliant III SYBR Green qPCR Master Mix with High ROX (7 μL) in a total volume of 15 μL. PCR conditions were 95 °C for 10 min, followed by 40 cycles of 95 °C for 30 s and 60 °C for 1 min. All qPCR reactions were normalized using the Ct value corresponding to the EiF4Ɣ peach gene (Pp_Eif4Ɣ) and the 60 S L13 ribosomal protein gene (Pp_RPL13). Each sample was measured three times with three technical replicates. The list of the designed primers for the qPCR is in **Table S2**.

### 2.4 Nucleic Acid Extraction, Illumina sequencing and Genotyping

Two resistant peach trees, Nemared and WFP, and two susceptible peach trees, Montclar and Pamirskii were used in this study. The genomic DNA of all tested plants was extracted using 100 mg of frozen young leaves. Each sample was ground with a mixer mill in 330 μL of extraction buffer (sorbitol 0.35 M, Tris 0.1 M, EDTA 5 mM, 4 mg of sodium metabisulfite), 330 μL of lysis buffer (Tris 0.2 M, EDTA 50 mM, NaCl 2 M, CTAB 2%), and 130 μL of sarkosyl 5%. After a chloroform-isoamyl alcohol (24:1) procedure, precipitation with isopropanol and three ethanol washes, the DNA content was eluted in water and treated with RNAse. PCRs were carried out using the GoTaq® G2 Flexi DNA Polymerase or the Expand long-range kit for longer fragments, following the manufacturer’s instructions. DNA libraries were prepared according to Illumina Nextera’s instructions and quantified using an Agilent high-sensitivity chip (Duval et al. 2022). 125 paired-end sequencing was performed using an Illumina Hiseq 2500, and data pre-processing and sample quality checks were carried out. Library preparations and sequencing were carried out by the MGX-Montpellier platform.

### 2.5 Bacterial Artificial Chromosome (BAC)

A BAC library was constructed for the resistant peach WFP using the approach described in (Duval et al. 2022) to obtain the region encompassing the RMia gene. The WFP BAC library was screened for the SP91 and SP92 region using 3 labeled probes, which were designed using the Primer 3 software (**Table S2**). Selected clones were sequenced using the PacBio RS II sequencer. After a demultiplexing step, the sequence assembly was performed following the HGAP PacBio workflow (Chin et al. 2013)

### 2.6. PacBio Technology-Based Full-Length cDNA Library Construction, Preparation and Sequencing

Total RNA was extracted from two nematode-infested root samples using the NucleoSpin® RNA Plant kit (Macherey Nagel). The quality of total RNA was evaluated using an Agilent 2100 Bioanalyzer (Agilent Technologies, CA, USA). Only samples with RNA integrity number (RIN) ≥ 8 were used for deep sequencing. To build the library for PacBio sequencing, RNA qualified was reverse transcribed using the SMARbell® Express Template Prep Kit 2.0. PacBio Sequel® Systems was used to sequence transcriptomes. The Sub-reads shorter than 50 bp, the poor-quality reads and specific adaptor sequences were removed (CTCACAGTCTGTGTGT for reverse primer of RNA roots). We used the Iso-Seq 3 bioinformatics workflow in SMRT Link Version 7.0.1.66975 to produce full-length transcript sequences and splice variants of good quality, with no assembly required. The Iso-Seq analysis workflow starts with the generation of high-fidelity reads using the circular consensus sequencing (CCS) method and the required trimmed poly(A) tail. The minimum accepted subread length was 50 bp, and we added the isoseq-mode option. We used the Iso-Seq 3 bioinformatics workflow in SMRT Link Version 7.0.1.66975 to produce full-length transcript sequences and splice variants of good quality, with no assembly required. The Iso-Seq analysis workflow starts with the generation of high-fidelity reads using the circular consensus sequencing (CCS) method and the required trimmed poly(A) tail. The minimum accepted subread length was 50 bp, and we added the isoseq-mode option.

### 2.7. sequence assembly and analysis

The sequence mapping on the peach genome V2 was obtained using BWA (v 0.7.12-r1039) with the BWA-MEM algorithm for mapping. The read alignment and index files (BAM and BAI files), genomic feature files (GFF3) on the reference Peach V2, and *de novo* sequences were visualized with the Integrative Genomics Viewer (Robinson et al. 2011). We used the QIAGEN CLC Genomics Workbench (https://digitalinsights.qiagen.com) for the sequence trimmings, gene annotations, and motif research. The software used for the gene prediction was FGENESH (Solovyev et al. 2006), with *P. persica* as organism-specific gene-finding parameters. Protein analysis and domain research were carried out using Interproscan (Paysan-Lafosse et al. 2023). Synteny was visualized using Kablammo software (Wintersinger and Wasmuth 2015)

## 3. Results

### Genomic region encompassing the RMia locus in the resistant varieties WFP and Nemared

A fine mapping study localized the *RMia* locus in the peach resistant variety ‘Nemared’, in the chromosome 2 between two KASP™ markers, SP92 and SP91 (Duval et al. 2014). In the first version of the peach reference genome (Lovell; v1.0) (Verde et al. 2013), four TNL genes were annotated in this interval, ppa023253m, ppa018595m, ppa000501m and ppa023503m (**Figure 1a**). In the next version of the reference genome (v2.0) (Verde et al. 2017), the region delineated by the same markers, contains 2 complete TNL genes (Prupe.2G055600 and Prupe.2G055700), 1 truncated TNL (Prupe.2G055500) (**Figure 1b**) and 3 other genes coding for, a trafficking protein particle complex, a short transmembrane protein and an unknown protein. Amplification with specific primers of these three TNL by PCR using Nemared DNA failed, suggesting that some polymorphism exists in the region. We therefore carried out NGS sequencing of both resistant accessions Nemared and WFP and the susceptible rootstocks Montclar and Pamirskii. While mapping NGS short reads of both resistant genotypes on the SP92-SP91 region of the peach reference sequence (v2.0), we observed that a large part of this region was not covered by reads, in line with the presence of a large indel in this region (**Figure S1ab**). Conversely, the same experiment using the reads of the susceptible rootstocks Montclar and Pamirskii led to a complete mapping onto the reference sequence meaning that no such large indels exist in both susceptible varieties. Consequently, we postulated that this targeted sequence was distinct between, on one hand, the susceptible rootstocks Lovell, Montclar and Pamirskii, and on the other hand, the resistant accessions WFP and Nemared. A BAC library was constructed to obtain the sequence of the SP92-SP91 region in the resistant genotype WFP. We designed specific probes inside this region to screen the BAC clones (**Table S2**). After aligning the BAC ends of five screened BACs on the Peach V2, we selected one BAC (43N01) and sequenced it using PacBio long-read sequencing technology. We confirmed the accuracy of the BAC sequence by mapping the WFP NGS short-reads on this *de novo* BAC sequence. In addition, we also mapped the NGS short-reads of Nemared on the BAC sequence and we observed that the SP92-SP91 regions are similar in Nemared and WFP (**Figure S1c**). This sequence, named hereafter ‘R_SP92SP91’, encompassing the *RMia* locus is 145198 kb long. Therefore, the R_SP92SP91 region is 71.7 kb larger than the V2_SP92-SP91 region in the peach reference genome (v2.0). The visualization of the syntenic region between both sequences indicated that this extended sequence is mainly the result of a large insertion in the central part of the region (**Figure S2**).

**Figure 1:**
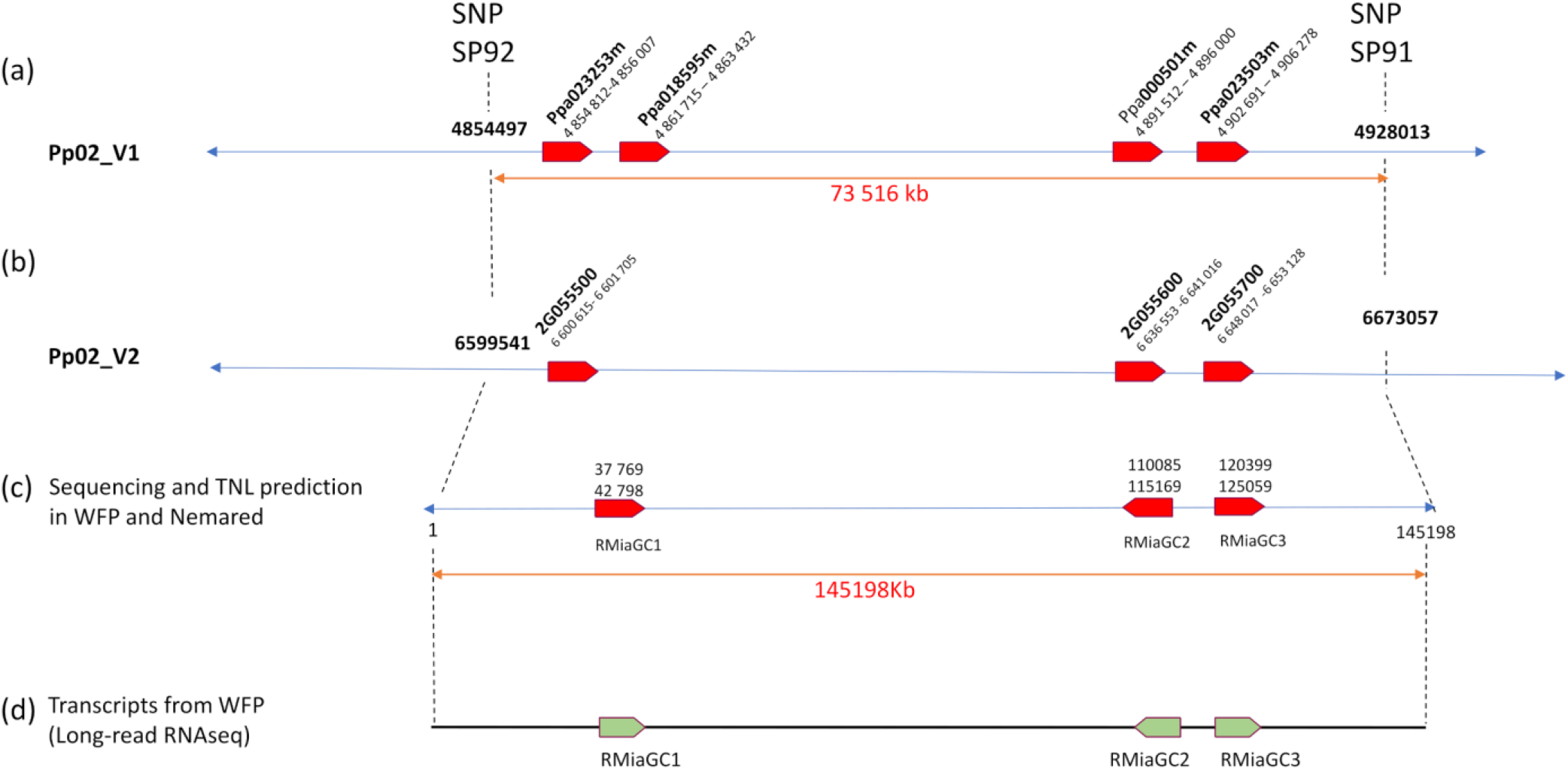
Schematic representation of the genomic interval ‘SP92-SP91’ from the chromosome 2. (a) TNL positions in the peach reference genome (*P. persica cv. Lovell*) V1. (b) TNL positions in the V2. (c) Position of predicted TNL genes in the resistant varieties Nemared and WFP. (d) Expressed genes in the SP92-SP91 interval.

### Structural and functional annotation of the R_SP92SP91 region

The analysis of the R_SP92SP91 sequence revealed that three full-length TNL genes (hereafter named as RMiaGC1, RMiaGC2 and RMiaGC3) were predicted along with 35 other putative genes including a large majority of genes coding for small peptides with no functional annotation and several transposable elements (**Table S3**). RMiaGC2 and RMiaGC3 are arranged only 5 kb apart from each other in a head-to-head organization while RMiaGC1 is located 67 kb upstream (**Figure 1c**). We did a long-read RNAseq analysis from RNA of nematode-infected roots to assess (i) the predicted structural annotation and (ii) the expression of the predicted genes in the region. Only the three TNL genes were expressed in the region R_SP92SP91 (**Figure 1d**). The structure of the expressed genes, including the number and location of exons, introns, and UTR, was ascertained by mapping the full-length transcripts on the *de novo* BAC sequence. While RMiaGC1 exhibits only one transcript (**Figure S1c**), RMiaGC2 and RMiaGC3 display 5 and 3 isoforms, respectively. As isoforms mainly differ in the 3’ and 5’ UTR length, it may affect the expression regulation of these candidates but unlikely their protein sequences. Only one isoform of the RMiaGC2 displays a slight variation in the LRR domain. RMiaCG1, RMiaGC2 and RMiaGC3 code for 3 predicted proteins of 1178 aa, 1154 aa, and 1095 aa, respectively (**Figure 2**). The percentage identity between these 3 genes ranged between 74.56% to 80.17% (**Table S4**). A BlastP search using the three candidate genes as queries against the proteome of Lovell, detects the highest identity for RMiaGC1 and RMiaGC2, (79% and 71%, respectively) with a TNL (Prupe.2G018300), located outside the SP92SP91 region, 4.88 Mb away on the chromosome 2 (**Table S4**). The percent identity between these 3 genes and the 3 TNL present in the SP92-SP91 region of the peach reference sequence (*i.e*. Prupe.2G055500, Prupe.2G055600 and Prupe.2G055700) ranged from 59.55% to 83.72% (**Table S4**). The functional annotation revealed the canonical organization of TNL proteins (*i.e*. the following chain of domains from the N-terminus to the C-terminus end: TIR, NB-ARC, NLL, LRR and PL-CJID) (**Figure 3b**). As expected, the 3 first domains exhibit a low polymorphism with all the canonical/functional motifs described in the literature. The LRR domain is also relatively conserved between the 3 candidates, except a short deletion of 10 aa located in RMiaGC2 (**Figure S3**). Conversely, PL-C-JID domains show a higher polymorphism, including larger indels. Beyond these discrepancies, the 3 PL-C-JID domains, harbor the classic motifs of the domain and atypical acidic regions made of repeat of aspartic and glutamic acids (D and E) (**Figure 3 and Figure S3)**. While polymorphic regions between motifs 1 and 2 and in the C-terminal end involve regions rich in acidic amino acids, polymorphic regions between motifs 2 and 4 are enriched in polar amino acids.

**Figure 2.**
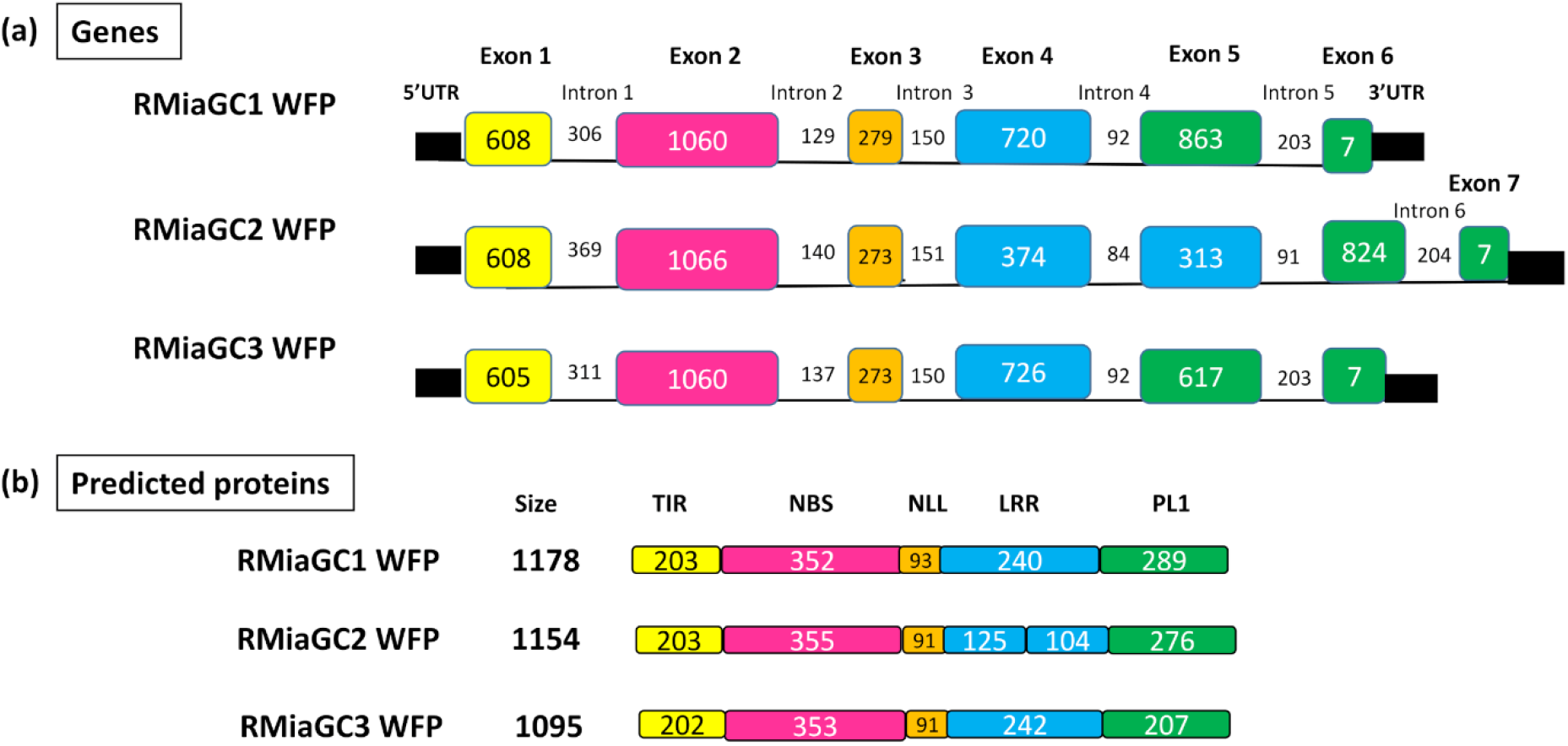
Structural annotation of the three RMia gene candidates. (a) Structure of the three genes RM1aCG1, RMiaGC2 and RMiaGC3. Size of exons and introns are in bp. Exons are represented in boxes, introns by black lanes. (b) Predicted domains of the three RMia candidate proteins in amino acids.

**Figure 3:**
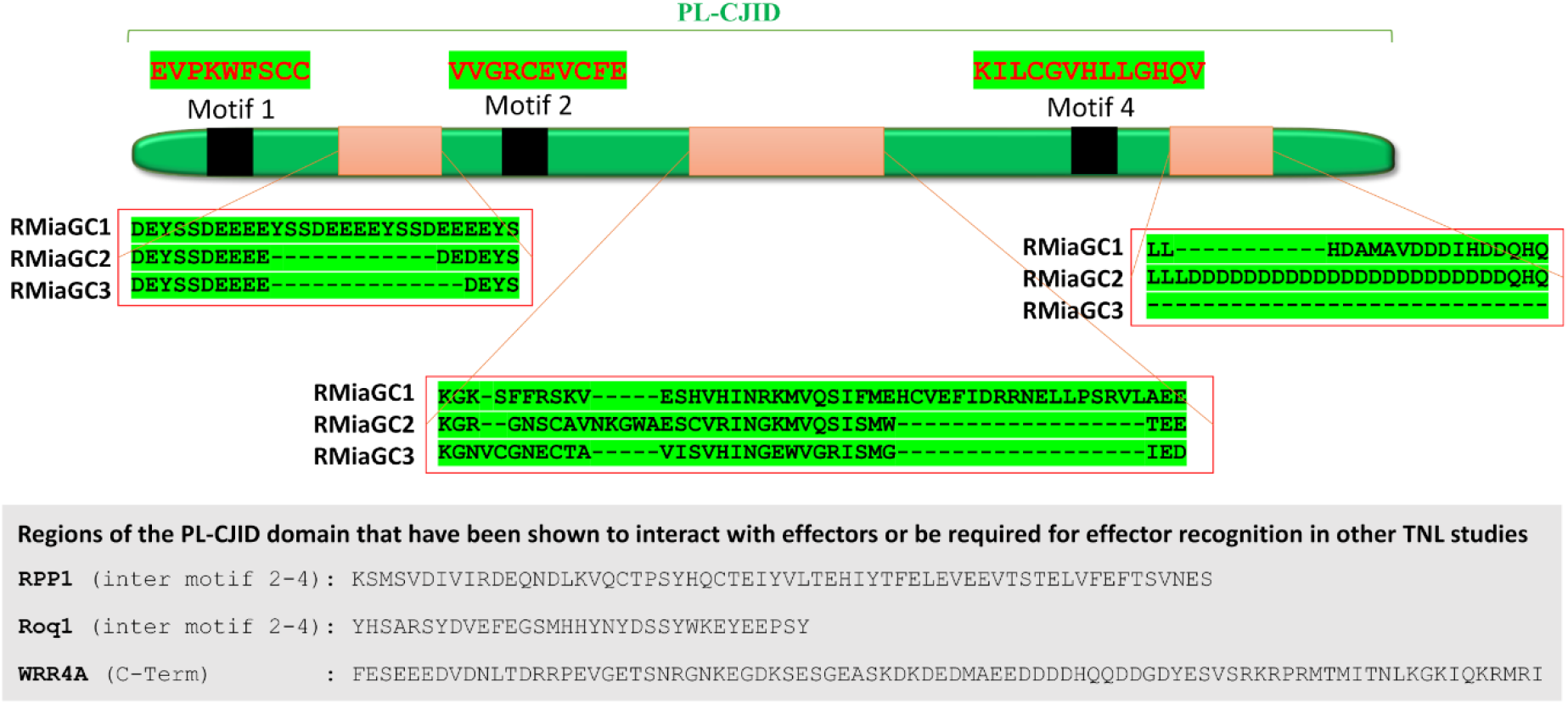
Detailed view of the PL-CJID domain of the three RMia gene candidates. The picture highlights the polymorphic regions between the three RMia candidates and the existing knowledge from other TNL studies for RPP1 (Ma et al. 2020), Roq1 (Martin et al., 2020) and WRR4A (Castel et al. 2019).

### Quantitative expression analysis of RMia candidate genes

We analyzed the relative normalized expression profiles of RM1aCG1, RMiaGC2 and RMiaGC3 from WFP seedlings infested with nematodes and from seedlings free of nematodes, using specific primers for each candidate gene (**Table S2**). The RMiaGC1 gene was statistically strongly up-regulated 10 days after nematode infection **(Figure 4)**. RMiaGC2 and RMiaGC3 were also statistically up-regulated upon nematode infection but in a clear restrained manner. RMiaGC2 and RMiaGC3 harbor a similar intensity in response to RKN infection. We repeated the experiment using a resistant peach hybrid, PRMN Z64P40, obtained from *P. persica* (Pamirskii × Rubira) × (Montclar × Nemared). In this crossing, the resistance trait to RKN is exclusively carried by Nemared, which contains the RMia gene, whereas the three other parents are fully susceptible to RKN. Interestingly, the expression pattern of the 3 candidate genes follows the same trend, thus confirming that a similar regulation occurs within this cluster of TNL in response to RKN.

**Figure 4:**
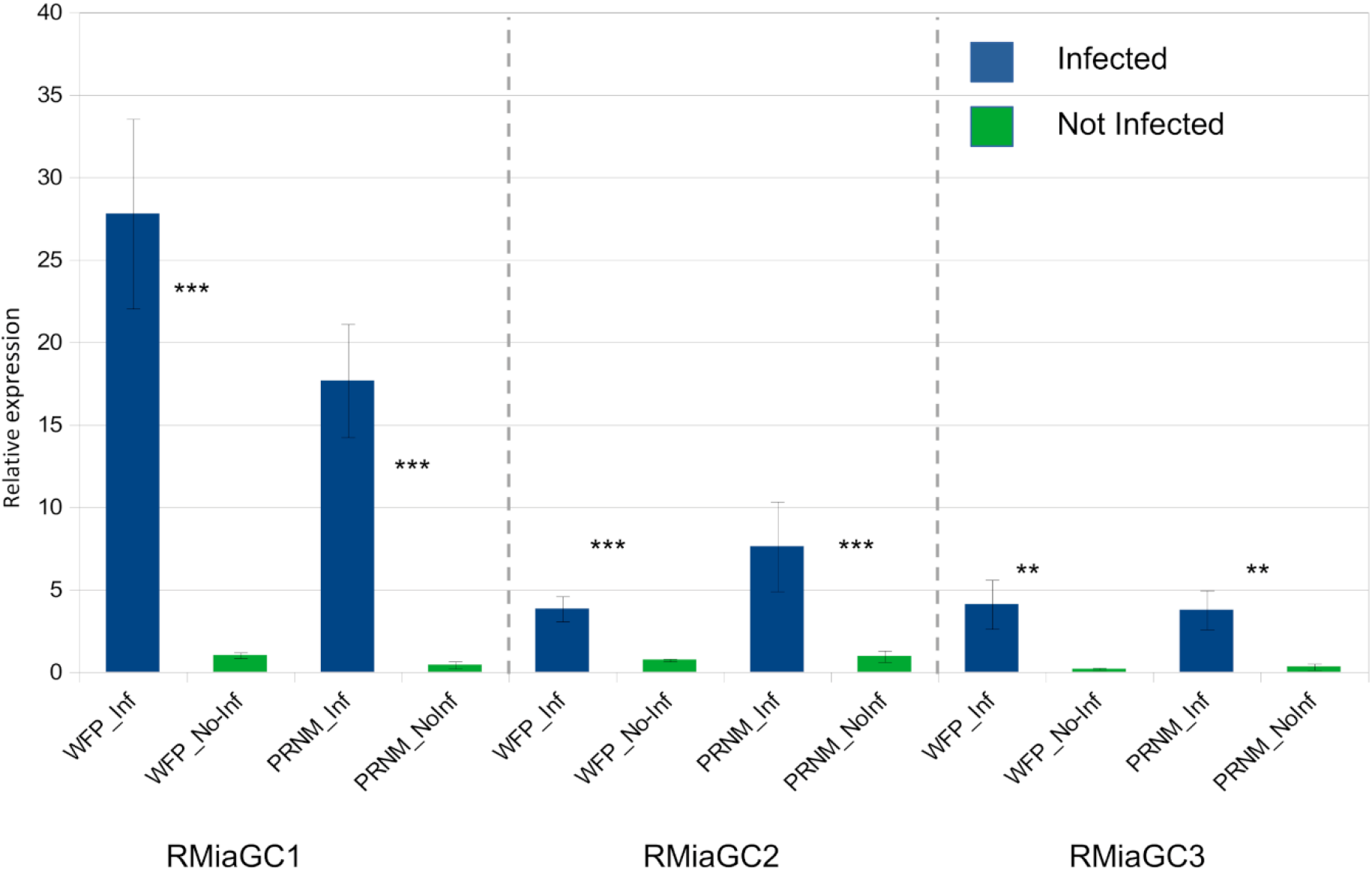
Expression pattern of the three genes RM1aCG1, RMiaGC2 and RMiaGC3 in the resistant WFP and PRMN hybrid seedlings in control (Noinf) and nematode-infected plants (Inf). Data represented the mean of three biological replicates. *** Significative differences between inoculated and control plants (_Inf vs. _Noinf). Differences in relative gene expression were statistically assessed using a Kruskal-Wallis test (***P<0.001, **P<0.01).

### Marker development for marker assisted selection

The acquisition of the full sequence of the resistant locus was an essential prerequisite to develop reliable molecular markers. We designed specific intragenic molecular markers in the genes RMiaGC1 and RMiaGC2 to cover the region (**Table S1**). The position of the markers used is represented in the **Figure 5a**. For the gene RMiaGC1, the two markers are G13F1R1 designed for the qPCR expression and another one in the 5’UTR region, G13QP211. For RMiaGC2, we designed a reliable marker, G13F5R5. The **Figure 5b** shows the results of the PCR amplifications for two resistant varieties (WFP and Nemared) with one band for the three markers and no amplification for three susceptible varieties (Montclar, Rubira and Pamirskii). A single signal was visible in the resistant varieties for the markers G13F1R1, G13QP211 and G13F5R5 at 437 bp, 150 bp and 205 bp, respectively. We tested these markers in other peach rootstocks for which RKN-related phenotypic status had been established. In all accessions, the genotype corresponded to the phenotype when using the molecular markers SP92, G13F1R1, G13QP211 and G31F5R5. Conversely, the KASP™ marker SP91 gave some discrepancies between genotypes and phenotypes for Nemaguard, the hybrid (Nemaguard x P1908), the peach-almond rootstocks Hansen 536, GF557 and the peach Missour 61-3, meaning that this marker was not tightly linked to the RMia resistance source (**Table 1**).

**Table 1:** Genotypes and phenotypes of resistant and susceptible peach rootstocks using the KASP™ markers SP92 and SP91, and the molecular markers G13F1R1, G13QP2111 and G31F5R5 designed in this study. *Presence of a second band.

| Rootstocks | Phenotype<br>vs. *M. incognita* | Markers |  |  |  |  |
| --- | --- | --- | --- | --- | --- | --- |
|  |  | SP92 | G13F1R1 | G13QP211 | G31F5R5 | SP91 |
| WFP Pleureur | R | RR | R | R | R | RR |
| Nemared | R | RR | R | R | R | RR |
| Montclar® | S | SS | S | S | S | SS |
| Rubira | S | SS | S | S | S | SS |
| Pamirskii | S | SS | S | S | S | SS |
| Nemaguard | R | RS | R | R | R | SS |
| (Nemaguard xP1908)5 | R | RS | R | R | R | SS |
| Hansen 536 | R | RS | R | R | R | SS |
| GF557 | R | RS | R* | R | R | RR |
| Garnem (GN15) | R | RS | R | R | R | RR |
| Summergrand | R | RR | R | R | R | SS |
| Missouri 61- 3 | R | RR | R | R | R | SS |
| GF305 | S | SS | S | S | S | SS |

**Figure 5:**
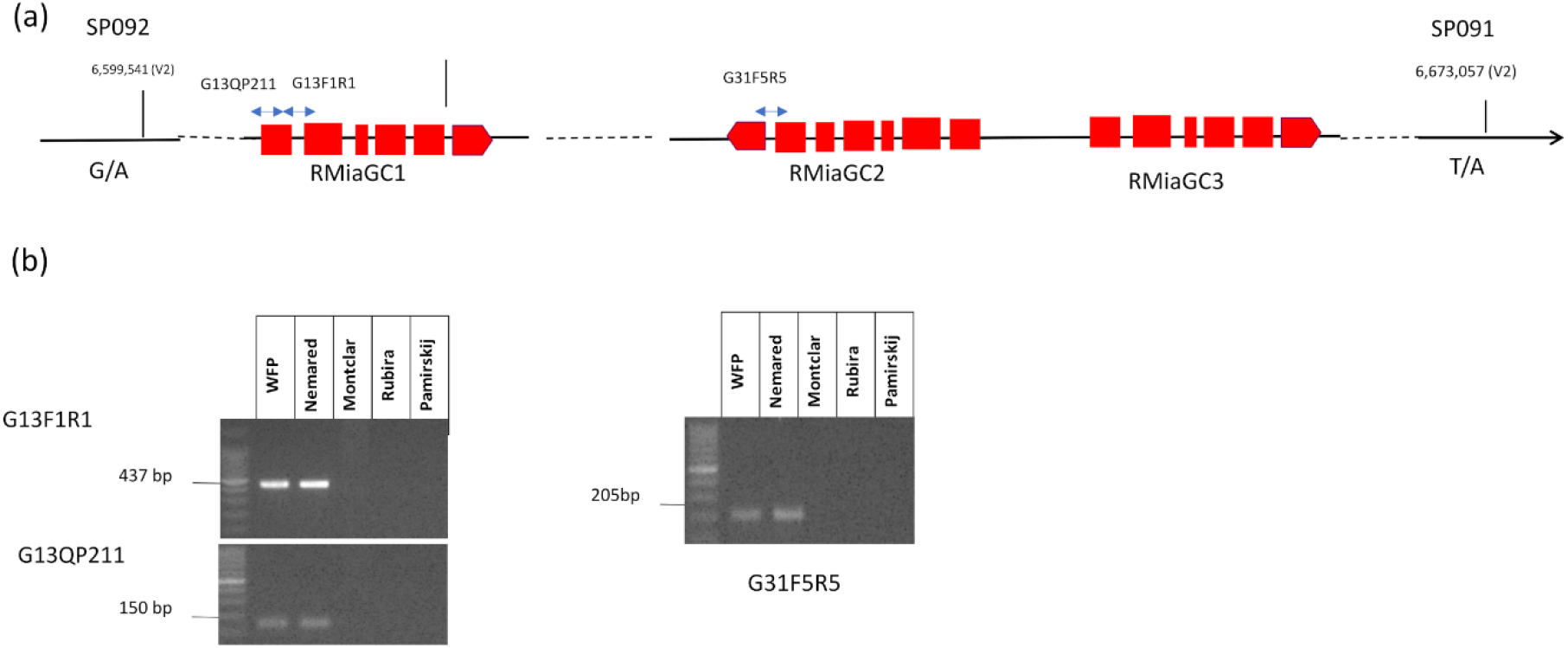
Genotyping using molecular markers designed in the RMia region for resistant and susceptible peaches. (a) Position of the framing markers SNP092 and SP091 and specific markers b) DNA amplifications for the RMiaGC1 and RMiaGC2 genes with the primers G13F1R1, G13QP211 and G31F5R5, respectively

## 4. Discussion

### Structural variation in the SP92SP91 region

In 2014, we identified the genomic location of the RMia gene in a region of the chromosome 2 marked out by two SNPs (SP92 and SP91), where four TNL genes were annotated. We observed that almost all susceptible varieties contained a retrotransposon inserted between the two truncated TNL genes, ppa023253 and ppa018595 (Duval et al. 2014). Considering their structure, we hypothesized that both genes initially formed a single functional gene, RMia, that might be inactivated due to the transposon’s insertion. However, we were unable to amplify the whole gene from the resistant peaches Nemared and WFP with specific primers flanking both genes ppa023253 and ppa018595m. In addition, we found other susceptible peach varieties such as Pamirskii, that did not include the transposable element. The results of the present study clearly indicate that other genes are present in WFP and Nemared compared to the peach reference genome. The origin of these genes could be either an introgression in this region or could originate from duplication followed by rearrangements in the region as we also found the highest percent identity between the first candidate gene, RMiaGC1, and Prupe.2G018300, a gene located upstream the SNP SP92 in the peach reference genome. However, considering the rather weak percentage identity between the 3 RMia candidates and the NLRome from the peach reference genome, the RMia gene likely originated from introgression coming from a source that still needs to be discovered. As Nemared and WFP are two varieties that are not directly genetically related, the introgression might have occurred some time ago in a common ancestor.

### Regulation of RMia candidate genes following RKN infection

The long-read RNAseq analysis obtained from WFP indicated that the three TNL genes, RMiaGC1, RMiaGC2 and RMiaGC3 were expressed in roots. The qPCR analysis showed that the RMiaGC1 gene is the most upregulated gene in the presence of nematodes. The overexpression of RMiaGC1 was confirmed by the same qPCR analysis in the other resistant genotype (PRMN140) including the resistant gene from Nemared. By contrast, the other two candidate genes (RMiaGC2 and RMiaGC3) studied in the same analysis are also overexpressed but at a lower level. Interestingly, both genes display a similar regulation when infected by nematodes that may be explained by shared regulatory elements, likely in the 5kb region between them. Similarly to other NLR genes (Borrelli et al. 2018), the three candidate genes are constitutively expressed at a low level and are upregulated, at different levels of expression, following pathogen’s infection. If pathogens usually dampen the immune system in susceptible plants leading to infection, in resistant plants, NLR are often upregulated directly or as a secondary consequence (Castel et al. 2019). The paradigm for NLR has recently evolved. Intricate networks between ETI and PTI components work in concert from the invader’s detection to activation of multiple NLR and formation of “resistosome” structures leading to cell death and resistance (Huang et al. 2023). Although sensor NLR are not systematically upregulated, the HR process and resistance come with accumulation of transcripts and proteins of activator/helper NLR (Borrelli et al. 2018). On one hand, the transcription of RMiaGC1 is strongly responsive to RKN infection. On the other hand RMiaGC2 and RMiaGC3 are localized in close proximity to each other and organized in head-to-head configuration that reminds the genetic and functional paired NLR (e.g. RGA4/RGA5 and RPS4/RRS1) (Narusaka et al. 2009; Cesari et al. 2014; Xi et al. 2022). In our study, the RMia resistance might be mediated by one of the candidate genes or the result of the cooperation of 2 or even the 3 candidate genes. One hypothesis may be that RMiaGC2 and RMiaGC3 act similarly to paired TNL (sensors/executors). In addition, this paired TNL complex may induce a strong expression of RMiaGC1 leading to the resistance. Alternatively, the RMiaGC1 gene may be the sole RMia gene and would be able to induce the plant’s immune response to nematodes by the large accumulation of its transcripts. A similar cluster of TNL, involved in resistance to the aphid *Myzus persicae*, has been recently identified in the same variety (WFP) (Duval et al. 2022). In such cases, high-resolution mapping studies would hardly settle these intricate models due to the considerable number of hybrids that should be produced to reveal discriminating recombinants. On the other side, transformation of susceptible peaches, with the different candidates to reveal the RMia gene, would be possible in theory. However, and despite the fact that production of *Prunus* hairy roots using *Agrobacterium rhizogenes* has recently made progress (Xu et al. 2020), the functional validation of the different constructions (seven different possible combinations), which are necessary to elucidate the RMia model, would be indubitably challenging in this plant model. The heterologous expression in a model plant such as *Arabidopsis* or *Tobacco* (Xiao et al. 2021) would be an alternative, but with no guarantee of success as other components, required for upstream or downstream signalization, might be absent in such plants. Ultimately, the use of ‘new genomic techniques’ (NGTs) may be the answer to edit such candidates making their functional characterization easier in the near future (Nerva et al. 2023).

### RMia among resistance genes to RKN in Prunus

In recent years, different sources of resistance to RKN have been identified among perennial plants (Saucet et al. 2016). Among woody plants, *Prunus* species hosts several resistance genes to RKN with different spectra of resistance to the most common species of the *Meloidogyne* genus. Apart from the RMia gene in *P. persica*, a succession of TNL genes have been identified in *P. sogdiana*, the wild myrobalan plum, that confer a resistance to *M. incognita* (Zhu et al. 2017; Xiao et al. 2021). In addition, the widely studied Ma gene in *P. cerasifera* confers a full resistance to all *Meloidogyne* species tested (Claverie et al. 2011; Duval et al. 2019). Besides, the Ma ortholog, RMja, in Almond does not control some *Meloidogyne* species such *M. incognita* or *M. floridensis* but it presents the same unique structure as the Ma gene (Van Ghelder et al. 2018; Duval et al. 2019). As RMia confers resistance to *M. incognita* but not to *M. javanica*, it is very complementary with the other RKN resistance genes. Considering the gene structure, Ma and RMja have an atypical repetition of 5 PL-CJID domains, a structure suspected to be involved in the RKN recognition (Van Ghelder et al. 2018). Our study revealed that RMia gene candidates have a more conventional structure including the canonical domains of TNL. Recently, the PL-CJID domain, which is present in the large majority of TNL (Van Ghelder and Esmenjaud 2016), has been shown to be directly involved in the pathogen recognition in different TNL models. Indeed, structural studies demonstrated that effectors interacted with polymorphic regions between conserved motifs of the PL-CJID domains of RPP1 (Ma et al. 2020) and Roq1 (Martin et al. 2020). Later, the extreme C-terminal end of WRR4A was shown to be essential for the effector recognition of the oomycete *Albugo candida* (Castel et al. 2021). In our study, we shed light on the polymorphism that exists in these specific regions of the PL-CJID domain between RMiaGC1, RMiaGC2, and RMiaGC3. These differences are made of indels and repeats of specific amino acids. Similarly to the PL-CJID domains of WRR4A, the RMiaGC2 C-terminal end is enriched in aspartic and glutamic acids. The three candidate genes also contain repeats of acidic amino acids between motif 1 and 2. Besides, the region between motif 2 and 4 is rich in polar residues. Such type of amino acids has been shown to be involved in effector recognition in RPP1 (amino acids N, S, Q, and T) and Roq1 (NR loop). These regions in the RMia gene candidates might be the seat of the RKN effector interaction and recognition. The functional validation of this hypothesis would require the knowledge of interacting effectors, however very rare models involving a R gene together with its cognate nematode effector are described in the literature.

### Durability of RMia and pyramiding of R genes

While the Ma gene is effective against all *Meloidogyne* species, the nematode species, *M. javanica, M. enterolobii, and M. floridensis* recently described in Florida (Handoo et al. 2004), are able to infest rootstocks carrying the RMia gene. More recently, Qiu and colleagues tested two isolates of *M. arenaria* that produce galls on the peach rootstock Flordaguard, whose source of resistance is Okinawa peach, in which the RMia gene is present (Qiu et al. 2021). Even though both Ma and RMja genes have not been circumvented, this study shows that some RKN may be able to overcome resistance genes in *Prunus*. INRAE’s Prunus rootstock breeding programs aim to combine the three main dominant resistance genes, Ma (plum), RMja (almond) and RMia (peach) into new rootstocks to increase the durability of nematode resistance. Marker-Assisted Selection (MAS) for the Ma gene is possible using intragenic SCAR markers, CT3-4N and NSCAFLP2 (Claverie et al. 2011). For the RMja gene, MAS selection can be carried out using a specific SNP on chromosome 7 at position 9019871 in the almond reference genome (*P. dulcis* cv ‘Texas’ V2), using either the KASP™ markers SP903 and SP904 (Duval et al. 2018), or SNPid 52023 from the Affyx almond array chip (Duval et al. 2023). Regarding RMia, we designed the novel markers G13F1R1, G13QP211, and G31F5R5 and used the KASP™ marker SP92 (Duval et al. 2014) usable in MAS. We developed two robust intragenic markers, G13F1R1 and GQP211, within the RMiaGC1 gene. Besides, the KASP™ marker SP92 is the closest intergenic marker to RMiaGC1 (37.8 Kb) and was shown to be a good indicator as well. For all genotypes tested, including varieties and hybrids, the genotyping using SP92, G13F1R1, G13QP211, and G31F5R5 markers strictly follows the phenotypes, which make them linked to the RMia resistance, unlike the KASP™ marker SP91 for which phenotypic predictions were not always correlated to the genotype (**Table 3**). In conclusion, *Prunus* rootstock breeders have effective markers at their disposal for pyramiding the three genes Ma, RMja and RMia, in a three-way rootstock either with size markers or SNP markers.

## Data availability statement

Raw data related to *P. persica* WFP can be retrieved in the BioProject number **PRJNA854365** including Illumina WGS of WFP (**SRR20115452**) and long read RNAseq of WFP (**SRR20225451**).

The R_SP92SP91 sequence containing the three RMia gene candidates from WFP can be retrieved from the GeneBank accession number **PQ432884**.

Raw data related to the WGS of *P. persica* Pamirskii (**SAMN43405486**), Montclar (**SAMN43416978**) and Nemared (**SAMN43416965**) are publicly available in BioProjects number **PRJNA1154452** and **PRJNA854381**.

## Author Contributions

CVG and HD conceptualized the study. CVG, LH and HD analyzed the sequences and wrote the manuscript. LH performed the qPCR experiments and analysis, the sequencing data analysis (mapping, assembly) and the sequence submission in NCBI. ND performed DNA extractions and PCR analyses, CCar performed the RKN production and identification. HD performed the nematode infection tests; CCal performed the BAC screening and the BAC sequencing and assembly. All authors revised and approved the manuscript for publication.

## Acknowledgments

The authors thank the genotoul bioinformatics platform Toulouse Occitanie (Bioinfo Genotoul, https://doi.org/10.15454/1.5572369328961167E12) (accessed on 1 January 2018) for providing computing, storage resources, the MGX Platform—Montpellier GenomiX for the NGS sequencing, the INRAE Gentyane platform for the long-read RNAseq sequencing and Jacques Lagnel for advice in the bioinformatic analysis. This study was supported by the INRAE Plant Biology and Breeding Division and the Plant – Nematode Interaction (IPN) team.

## References

Abad, P., Gouzy, J., Aury, J.-M., Castagnone-Sereno, P., Danchin, E.G.J., Deleury, E., Perfus-Barbeoch, L., Anthouard, V., Artiguenave, F., Blok, V.C., Caillaud, M.-C., Coutinho, P.M., Dasilva, C., De Luca, F., Deau, F., Esquibet, M., Flutre, T., Goldstone, J.V., Hamamouch, N., Hewezi, T., Jaillon, O., Jubin, C., Leonetti, P., Magliano, M., Maier, T.R., Markov, G.V., McVeigh, P., Pesole, G., Poulain, J., Robinson-Rechavi, M., Sallet, E., Segurens, B., Steinbach, D., Tytgat, T., Ugarte, E., van Ghelder, C., Veronico, P., Baum, T.J., Blaxter, M., Bleve-Zacheo, T., Davis, E.L., Ewbank, J.J., Favery, B., Grenier, E., Henrissat, B., Jones, J.T., Laudet, V., Maule, A.G., Quesneville, H., Rosso, M.-N., Schiex, T., Smant, G., Weissenbach, J., and Wincker, P. 2008. Genome sequence of the metazoan plant-parasitic nematode Meloidogyne incognita. Nat. Biotechnol. 26(8): 909–915. Nature Publishing Group, New York. doi:10.1038/nbt.1482.

Borrelli, G.M., Mazzucotelli, E., Marone, D., Crosatti, C., Michelotti, V., Valè, G., and Mastrangelo, A.M. 2018. Regulation and Evolution of NLR Genes: A Close Interconnection for Plant Immunity. Int. J. Mol. Sci. 19(6): 1662. doi:10.3390/ijms19061662.

Castel, B., Fairhead, S., Furzer, O.J., Redkar, A., Wang, S., Cevik, V., Holub, E.B., and Jones, J.D.G. 2021. Evolutionary trade-offs at the Arabidopsis WRR4A resistance locus underpin alternate Albugo candida race recognition specificities. Plant J. Cell Mol. Biol. 107(5): 1490–1502. doi:10.1111/tpj.15396.

Castel, B., Ngou, P.-M., Cevik, V., Redkar, A., Kim, D.-S., Yang, Y., Ding, P., and Jones, J.D.G. 2019. Diverse NLR immune receptors activate defence via the RPW8-NLR NRG1. New Phytol. 222(2): 966–980. doi:10.1111/nph.15659.

Cesari, S., Bernoux, M., Moncuquet, P., Kroj, T., and Dodds, P. 2014. A novel conserved mechanism for plant NLR protein pairs: the “integrated decoy” hypothesis. Front. PLANT Sci. 5. doi:10.3389/fpls.2014.00606.

Chin, C., Alexander, D., Marks, P., Klammer, A., Drake, J., Heiner, C., Clum, A., Copeland, A., Huddleston, J., Eichler, E., Turner, S., and Korlach, J. 2013. Nonhybrid, finished microbial genome assemblies from long-read SMRT sequencing data. Nat. Methods 10(6): 563-+. doi:10.1038/NMETH.2474.

Claverie, M., Dirlewanger, E., Bosselut, N., Van Ghelder, C., Voisin, R., Kleinhentz, M., Lafargue, B., Abad, P., Rosso, M.-N., Chalhoub, B., and Esmenjaud, D. 2011. The Ma Gene for Complete-Spectrum Resistance to Meloidogyne Species in Prunus Is a TNL with a Huge Repeated C-Terminal Post-LRR Region. Plant Physiol. 156(2): 779–792. doi:10.1104/pp.111.176230.

Djian-Caporalino, C., Navarrete, M., Fazari, A., Baily-Bechet, M., Marteu, N., Dufils, A., Tchamitchian, M., Lefèvre, A., Pares, L., Mateille, T., Tavoillot, J., Palloix (‡), A., Sage-Palloix, A.-M., Védie, H., Goillon, C., and Castagnone-Sereno, P. 2019. Conception et évaluation de systèmes de culture maraîchers méditerranéens innovants pour gérer les nématodes à galles. BASE. doi:10.25518/1780-4507.17725.

Duval, H., Coindre, E., Ramos-Onsins, S.E., Alexiou, K.G., Rubio-Cabetas, M.J., Martínez-García, P.J., Wirthensohn, M., Dhingra, A., Samarina, A., and Arús, P. 2023. Development and Evaluation of an AxiomTM 60K SNP Array for Almond (Prunus dulcis). Plants 12(2): 242. doi:10.3390/plants12020242.

Duval, H., Heurtevin, L., Dlalah, N., Callot, C., and Lagnel, J. 2022. The Rm1 and Rm2 Resistance Genes to Green Peach Aphid (Myzus persicae) Encode the Same TNL Proteins in Peach (Prunus persica L.). Genes 13(8): 1489. doi:10.3390/genes13081489.

Duval, H., Hoerter, M., Polidori, J., Confolent, C., Masse, M., Moretti, A., Van Ghelder, C., and Esmenjaud, D. 2014. High-resolution mapping of the RMia gene for resistance to root-knot nematodes in peach. Tree Genet. Genomes 10(2): 297–306. doi:10.1007/s11295-013-0683-z.

Duval, H., Van Ghelder, C., Callot, C., and Esmenjaud, D. 2018. Characterization of the RMja resistance gene to root-knot nematodes from the ‘Alnem’ almond rootstock. Acta Hortic. (1219): 325–330. doi:10.17660/ActaHortic.2018.1219.49.

Duval, H., Van Ghelder, C., Portier, U., Confolent, C., Meza, P., and Esmenjaud, D. 2019. New Data Completing the Spectrum of the Ma, RMia, and RMja Genes for Resistance to Root-Knot Nematodes (Meloidogyne spp.) in Prunus. Phytopathology 109(4): 615–622. doi:10.1094/PHYTO-05-18-0173-R.

Handoo, Z.A., Nyczepir, A.P., Esmenjaud, D., van der Beek, J.G., Castagnone-Sereno, P., Carta, L.K., Skantar, A.M., and Higgins, J.A. 2004. Morphological, molecular, and differential-host characterization of Meloidogyne floridensis n. sp (Nematoda: Meloidogynidae), a root-knot nematode parasitizing peach in Florida. J. Nematol. 36(1): 20–35. Soc Nematologists, Marceline.

Huang, S., Jia, A., Ma, S., Sun, Y., Chang, X., Han, Z., and Chai, J. 2023. NLR signaling in plants: from resistosomes to second messengers. Trends Biochem. Sci. 48(9): 776–787. Elsevier. doi:10.1016/j.tibs.2023.06.002.

Jones, J., Haegeman, A., Danchin, E., Gaur, H., Helder, J., Jones, M., Kikuchi, T., Manzanilla-Lopez, R., Palomares-Rius, J., Wesemael, W., and Perry, R. 2013. Top 10 plant-parasitic nematodes in molecular plant pathology. Mol. PLANT Pathol. 14(9): 946–961. doi:10.1111/mpp.12057.

Khallouk, S., Voisin, R., Van Ghelder, C., Engler, G., Amiri, S., and Esmenjaud, D. 2011. Histological Mechanisms of the Resistance Conferred by the Ma Gene Against Meloidogyne incognita in Prunus spp. Phytopathology 101(8): 945–951. doi:10.1094/PHYTO-01-11-0004.

Ma, S., Lapin, D., Liu, L., Sun, Y., Song, W., Zhang, X., Logemann, E., Yu, D., Wang, J., Jirschitzka, J., Han, Z., Schulze-Lefert, P., Parker, J.E., and Chai, J. 2020. Direct pathogen-induced assembly of an NLR immune receptor complex to form a holoenzyme. Science 370(6521): eabe3069. American Association for the Advancement of Science. doi:10.1126/science.abe3069.

Martin, R., Qi, T., Zhang, H., Liu, F., King, M., Toth, C., Nogales, E., and Staskawicz, B.J. 2020. Structure of the activated ROQ1 resistosome directly recognizing the pathogen effector XopQ. Science 370(6521): eabd9993. American Association for the Advancement of Science. doi:10.1126/science.abd9993.

Massonie, G., and Maison, P. 1979. Preliminary-Results on the Study of the Resistance of Prunus-Persica L (batsch) Varieties to Myzus-Persicae Sulz and Myzus-Varians Davids. Ann. Zool. Ecol. Anim. 11(3): 479–485. Inst Natl Recherche Agronomique, Paris Cedex 07.

Narusaka, M., Shirasu, K., Noutoshi, Y., Kubo, Y., Shiraishi, T., Iwabuchi, M., and Narusaka, Y. 2009. RRS1 and RPS4 provide a dual Resistance-gene system against fungal and bacterial pathogens. Plant J. 60(2): 218– 226. doi:10.1111/j.1365-313X.2009.03949.x.

Nerva, L., Dalla Costa, L., Ciacciulli, A., Sabbadini, S., Pavese, V., Dondini, L., Vendramin, E., Caboni, E., Perrone, I., Moglia, A., Zenoni, S., Michelotti, V., Micali, S., La Malfa, S., Gentile, A., Tartarini, S., Mezzetti, B., Botta, R., Verde, I., Velasco, R., Malnoy, M.A., and Licciardello, C. 2023. The Role of Italy in the Use of Advanced Plant Genomic Techniques on Fruit Trees: State of the Art and Future Perspectives. Int. J. Mol. Sci. 24(2): 977. doi:10.3390/ijms24020977.

Paysan-Lafosse, T., Blum, M., Chuguransky, S., Grego, T., Pinto, B.L., Salazar, G.A., Bileschi, M.L., Bork, P., Bridge, A., Colwell, L., Gough, J., Haft, D.H., Letunić, I., Marchler-Bauer, A., Mi, H., Natale, D.A., Orengo, C.A., Pandurangan, A.P., Rivoire, C., Sigrist, C.J.A., Sillitoe, I., Thanki, N., Thomas, P.D., Tosatto, S.C.E., Wu, C.H., and Bateman, A. 2023. InterPro in 2022. Nucleic Acids Res. 51(D1): D418–D427. doi:10.1093/nar/gkac993.

Qiu, S., Maquilan, M.A.D., Chaparro, J.X., Brito, J.A., Beckman, T.G., and Dickson, D.W. 2021. Susceptibility of Flordaguard peach rootstock to a resistant-breaking population of Meloidogyne floridensis and two populations of Meloidogyne arenaria. J. Nematol. 53: e2021–111. doi:10.21307/jofnem-2021-111.

Ramming, D., and Tanner, O. 1983. Nemared Peach Rootstock. Hortscience 18(3): 376–376.

Robinson, J.T., Thorvaldsdóttir, H., Winckler, W., Guttman, M., Lander, E.S., Getz, G., and Mesirov, J.P. 2011. Integrative genomics viewer. Nat. Biotechnol. 29(1): 24–26. Nature Publishing Group. doi:10.1038/nbt.1754.

Saucet, D., S.B.;. Van Ghelder, C.;. Abad, P.;. Duval, H.;. Esmenjaud. 2016. Resistance to root-knot nematodes Meloidogyne spp. in woody plants. New Phytol. 211(1): 41–56.

Shao, Z., Zhang, Y., Hang, Y., Xue, J., Zhou, G., Wu, P., Wu, X., Wu, X., Wang, Q., Wang, B., and Chen, J. 2014. Long-Term Evolution of Nucleotide-Binding Site-Leucine-Rich Repeat Genes: Understanding Gained from and beyond the Legume Family. Plant Physiol. 166(1): 217–234. doi:10.1104/pp.114.243626.

Solovyev, V., Kosarev, P., Seledsov, I., and Vorobyev, D. 2006. Automatic annotation of eukaryotic genes, pseudogenes and promoters. Genome Biol. 7(1): S10. doi:10.1186/gb-2006-7-s1-s10.

Van Ghelder, C., and Esmenjaud, D. 2016. TNL genes in peach: insights into the post-LRR domain. Bmc Genomics 17: 317. doi:10.1186/s12864-016-2635-0.

Van Ghelder, C., Esmenjaud, D., Callot, C., Dubois, E., Mazier, M., and Duval, H. 2018. Ma Orthologous Genes in Prunus spp. Shed Light on a Noteworthy NBS-LRR Cluster Conferring Differential Resistance to Root-Knot Nematodes. Front. Plant Sci. 9: 1269. doi:10.3389/fpls.2018.01269.

Verde, I., Abbott, A., Scalabrin, S., Jung, S., Shu, S., Marroni, F., Zhebentyayeva, T., Dettori, M., Grimwood, J., Cattonaro, F., Zuccolo, A., Rossini, L., Jenkins, J., Vendramin, E., Meisel, L., Decroocq, V., Sosinski, B., Prochnik, S., Mitros, T., Policriti, A., Cipriani, G., Dondini, L., Ficklin, S., Goodstein, D., Xuan, P., Del Fabbro, C., Aramini, V., Copetti, D., Gonzalez, S., Horner, D., Falchi, R., Lucas, S., Mica, E., Maldonado, J., Lazzari, B., Bielenberg, D., Pirona, R., Miculan, M., Barakat, A., Testolin, R., Stella, A., Tartarini, S., Tonutti, P., Arus, P., Orellana, A., Wells, C., Main, D., Vizzotto, G., Silva, H., Salamini, F., Schmutz, J., Morgante, M., Rokhsar, D., and Int Peach Genome Initiative. 2013. The high-quality draft genome of peach (Prunus persica) identifies unique patterns of genetic diversity, domestication and genome evolution. Nat. Genet. 45(5): 487–U47. doi:10.1038/ng.2586.

Verde, I., Jenkins, J., Dondini, L., Micali, S., Pagliarani, G., Vendramin, E., Paris, R., Aramini, V., Gazza, L., Rossini, L., Bassi, D., Troggio, M., Shu, S., Grimwood, J., Tartarini, S., Dettori, M.T., and Schmutz, J. 2017. The Peach v2.0 release: high-resolution linkage mapping and deep resequencing improve chromosome-scale assembly and contiguity. Bmc Genomics 18: 225. doi:10.1186/s12864-017-3606-9.

Wintersinger, J.A., and Wasmuth, J.D. 2015. Kablammo: an interactive, web-based BLAST results visualizer. Bioinformatics 31(8): 1305–1306. doi:10.1093/bioinformatics/btu808.

Xi, Y., Cesari, S., and Kroj, T. 2022. Insight into the structure and molecular mode of action of plant paired NLR immune receptors. Essays Biochem. 66(5): 513–526. doi:10.1042/EBC20210079.

Xiao, K., Zhu, H., Zhu, X., Liu, Z., Wang, Y., Pu, W., Guan, P., and Hu, J. 2021. Overexpression of PsoRPM3, an NBS-LRR gene isolated from myrobalan plum, confers resistance to Meloidogyne incognita in tobacco. Plant Mol. Biol. 107(3): 129–146. doi:10.1007/s11103-021-01185-1.

Xu, S., Lai, E., Zhao, L., Cai, Y., Ogutu, C., Cherono, S., Han, Y., and Zheng, B. 2020. Development of a fast and efficient root transgenic system for functional genomics and genetic engineering in peach. Sci. Rep. 10(1): 2836. Nature Publishing Group. doi:10.1038/s41598-020-59626-8.

Zhu, X., Xiao, K., Cui, H., and Hu, J. 2017. Overexpression of the Prunus sogdiana NBS-LRR Subgroup Gene PsoRPM2 Promotes Resistance to the Root-Knot Nematode Meloidogyne incognita in Tobacco. Front. Microbiol. 8. Frontiers. doi:10.3389/fmicb.2017.02113.

