## Supplementary material for "Identification and Expression of the peach TNL *RMia* genes for the Resistance to the Root-knot Nematode *Meloidogyne incognita*": Suplementary figures S1-3

Supplementary figures S1, S2 and S3

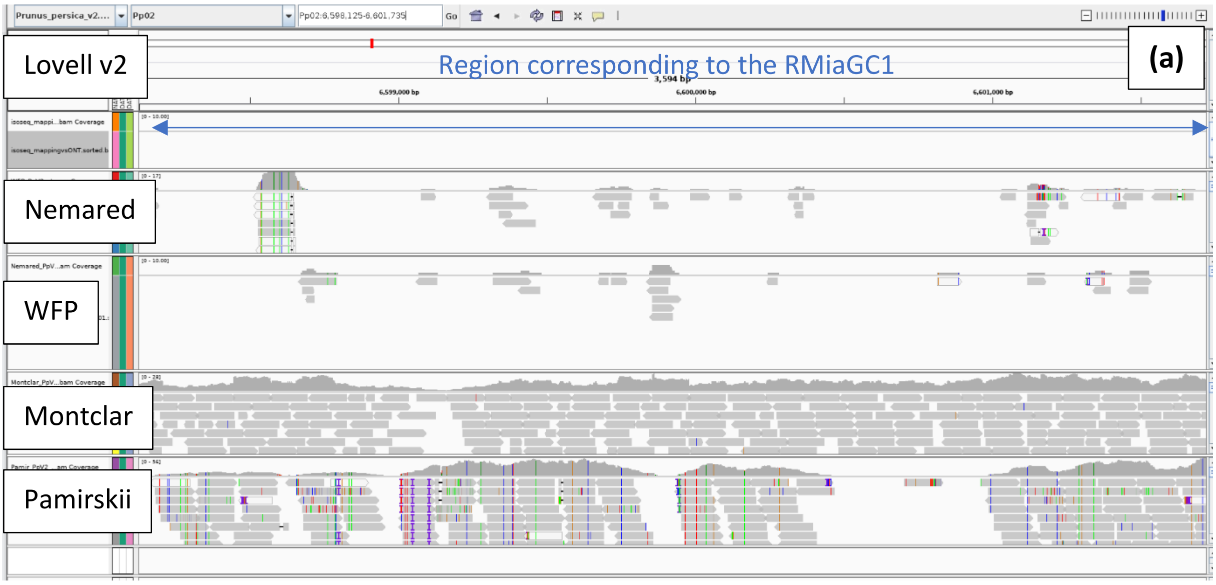


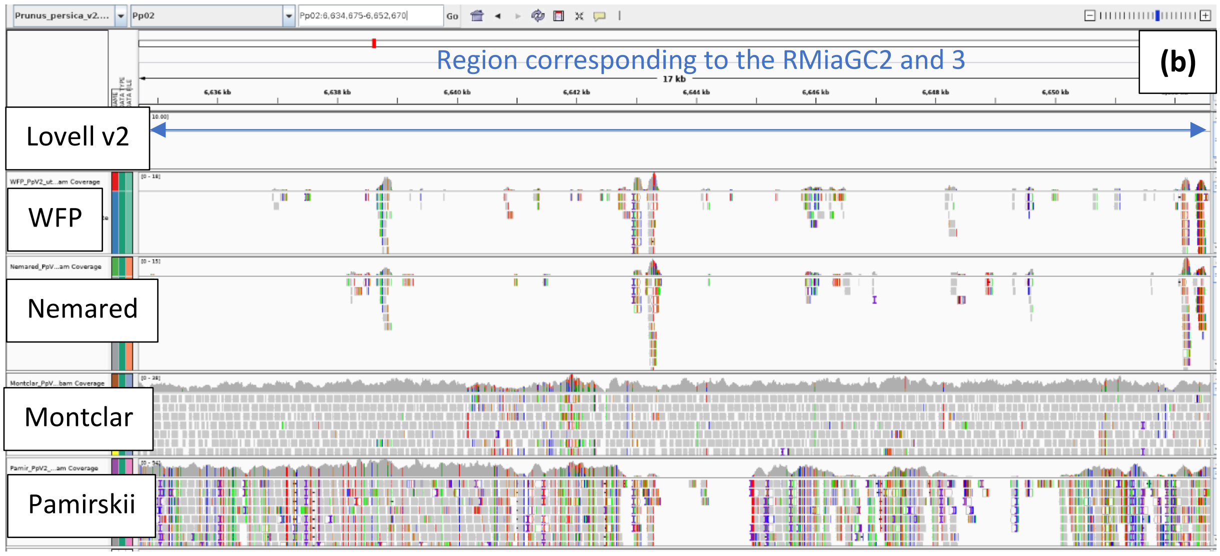

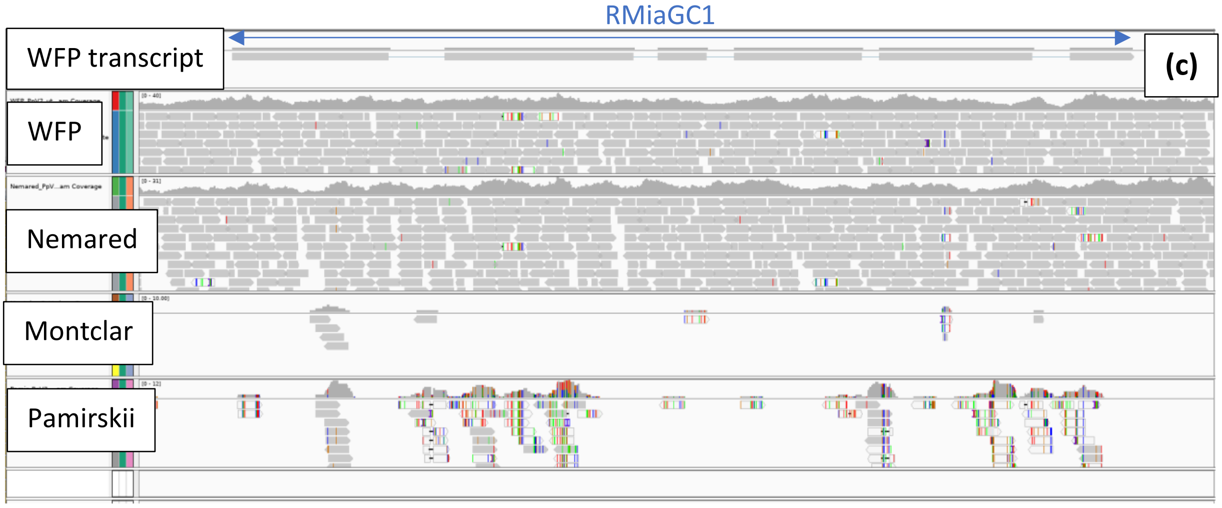


**Figure S1**: Mapping of the reads obtained from two resistant and two susceptible varieties of *P. persica* (WFP, Nemared and Montclar, Pamirskii) on regions corresponding to the 3 RMia gene candidates. Images are obtained using IGV software. (a) reads mapping of *P. persica* cv. WFP, Nemared, Montclar and Pamirskii on the reference peach genome v2 (Lovell). The region displayed correspond to the RMiaGC1 locus. (b) The same mapping corresponding to the region including RMiaGC2 and RMiaGC3 loci. (c) view of the alignment of the long-read RNAseq from WFP (WFP transcript), WFP, Nemared, Montclar and Pamirski. The region corresponds to the RMiaGC1 locus.

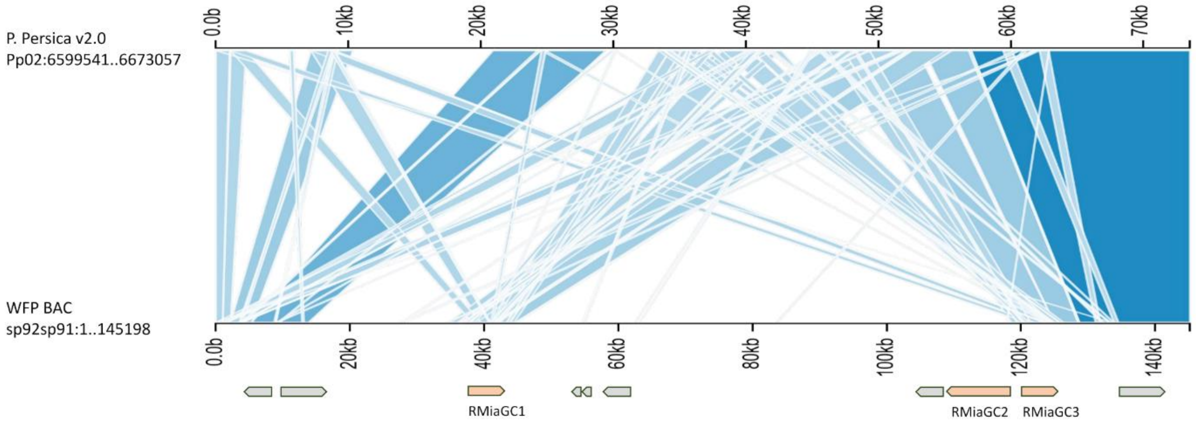


**Figure S2:** Synteny visualisation between the *P. persica cv. WFP* region sp92-sp91 and the corresponding region in the *P. persica* reference genome. The three RMia gene candidates (orange) are positioned together with the transposable elements (grey).

```
RMiaGC2         MFSLKSLFRLMGSSPLPIDSGVEPQLPSDLPPTTTQPVSSSPFSSSFAHDSSWTYDVFLS
RMiaGC3         MFSLKSLFRLMGSSPLPIDSGVEPQLPSDLPPTTTQPVSSSPSSSSFAHDSSWTYDVFLS
RMiaGC1         MFSLKSLFRLMGSSPLPIDSRVEPQLPSDLPPTTTQPVSSSPSSSSFAPDSSWTYDVFLS
                ******************** ********************* ***** ***********

RMiaGC2         FRGEDTRTNFTDHLYKALCDKGIYTFIDRELVRGEEISPALVKAIEESRISVIVFSENYA
RMiaGC3         FRGEDTRTNFTDHLYKALCDKGIYTFIDRELVRGEEISPALVKAIEESRIFVIVFSENYA
RMiaGC1         FRGEDTRTNFTDHLYKALCDKGIYTFIDRELVRGEEISPALVKAIEESRISLIVFSENYA
                ************************************************** :********

RMiaGC2         SSRWCLDELVKILQCKESKQQIVLPIFYKVDPSDVGNQKNRFGDAFKGLIERKFKNDKQK
RMiaGC3         SSRWCLDELVKILQCKESKQQIVLPIFYKVDPSDVRHQRSSYGDAFVHH-ERKFKDDREK
RMiaGC1         SSRWCLDELVKILQCKESKQQIVLPIFYKVDPSDVGNQKNRFGDAFKGLIERKFKNDKQK
                *********************************** :*.. :****    *****:*.:*

RMiaGC2         LLIWRKALEKAAKLSGYPSKDGEYETIFINNIVDGILNQVLSRTYLNVPKYPVGIQSHVQ
RMiaGC3         VLKWRRALKEAANLSGWHFEEGAYEAIFVNNIVDEIISRVLRRTYWNVAKHPVGIQSHVQ
RMiaGC1         LLIWRKALEKAAKLSGYPSKDGEYEAIFINNIVDEIISRVLRRTYWNVAEHPVGIQPHVQ
                :* **.**::**:***:  ::* **:**:***** *:..** *** **.::*****.***

RMiaGC2         DVEMLLDVGGNGRHMVGIWGTSGIGKTTIAKAICNAVGYKFEGSCFLPNVREGSLIQLQE
RMiaGC3         DVKKLLDVDGNGRRMIGIWGTSGIGKTTIAKAIWNAIAHEFEGSCFLPNVREGCLVQLQE
RMiaGC1         DVKRLLDVGGNGRRMVGIWGTSGIGKTTIAKAIWNAIAHEFEGSCFLENVREGSLVQLQK
                **: ****.****.*:***************** **:.::******* *****.*:***:

RMiaGC2         TLLQEILGVKEFKIASADKGISIIHKLLRHKKILLILDDVNQLEQLDNLAGVGWFGEGSR
RMiaGC3         TLLDKILG-KNLKIQSVDEGIGVIKKRLRHKKILLILDDVDHLEQLENLAGDDWFGEGSR
RMiaGC1         TLLHKYLG-KKLEIQSVDEGIGVIKERLRHKRILLILDDVDQLEQLKKLGGDDWFGEGSR
                *** : ** *:::* *.*:**.:*:: ****.********::****.:*.* .*******

RMiaGC2         VIITTQDRGLLKSHGIKLIYEVQKLCDCQALELFSLNAFGINEPPNDYLELASRAIAFAD
RMiaGC3         VIITTKNRRLLENREIELVYEVKKLDYNQALELFSWHAFRRSEPPEDYLELAQCAITFAG
RMiaGC1         VIITTKNRRLLENRKIELIYEVKKLDYNQALELFSWHAFRRSEPPEDYLELAQRAIAFAD
                *****::* **:.. *:*:***:**   ******* :**  .***:******. **:**.

RMiaGC2         GHPLALTILGRHLHNRDKTFWQVILDSFKGEPYTHIERILQKSYDALDDYAKEAFLDIAC
RMiaGC3         GLPLALTILGAHLRGIDILRWKDILKDYKGEPYTYIERILQKSYDALDHRAKEYFLDIAC
RMiaGC1         GLPLALTILGSHLRGIDIRRWQVILDGYEGEPYTHIERILQKSYDALDPCAKEYFLDIAC
                * ******** **.. *   *: **..::*****:*************  *** ******

RMiaGC2         FFNDEEKDYVLQIVPKSCIEVLVDKAMITIEWNYSILMHGLLANLGKDIVHKESLNDPGR
RMiaGC3         FFKGRYKDYVLQIVPPKVIEEFVDKALITIEGG-DILMHDLLANLGKDIVHKESPDDPGQ
RMiaGC1         FFKGEKKDYVPQVVPQKFIEEFRDKALINIEWS-KIVMHDLLANLGKDIVRKESPNDPGQ
                **:.  **** *:** . ** : ***:*.** . .*:**.**********.*** :***.

RMiaGC2         RSRLWLYEDVKRVLTENTGTRNIKGIMVKLQKPVEITLNPECFRNMINLQIFINHNASLC
RMiaGC3         RSRLWFYEDVIQVLTDCTGKRKIKGIMVKLPEPAEITLNPECFRKMVNLQIFISHNASLC
RMiaGC1         RSRLWFHEDVIQVLMESTGTRNIKGITVKLPEPAEITLNPECFRNMVNLQIFINHNASLC
                *****::*** .** : **.*:**** *** :*.**********:*:******.******

RMiaGC2         GHINYLPDALRLINWDRCQLQSLPPNFQGNCLVGFNMLRSHIRQLEG--FKHLPNLTYMK
RMiaGC3         GHINYLPNALRIIDWPSCQLQSLPPNFQGNCLVEFKMPGSHIRQLER--FKHVPNLTCMN
RMiaGC1         GHINYLPNALRLINWDRCQLQSWPPNFQGNHLVEFSMPRSHIRQLERFNFKLMPNLTSMN
                *******:***:*:*  ***** ******* ** *.*  *******   ** :**** *:

RMiaGC2         LEDCQFLEKIPDLSRIPNIKYLHIEKCTSLVEVDDSVGFLDKLVELIVYKCGKLMRFGTT
RMiaGC3         LRGCQFLEKIPDLSGIPNIKYLFLSDCTSLVSLDDSVGFLDKLVILDLGGCVNLTKFGRR
RMiaGC1         LSDCQSLEKIPDLSGIPNIKYLHLSNCTSLVEVDDSVGFLDKLLELHLDGCIKLTRFGTR
                * .** ******** *******.:..*****.:**********: * :  * :* .**

RMiaGC2         VGLKSLDTLSVSDCERLESFPE----------LQLEGNGIRELPLSTLELSGLRLLDNGG
RMiaGC3         LKLKSLETLYLKDCESLESLPEIEVKMESLRRLDMEGSGIRELPPSIKHLTGLRDLSLER
RMiaGC1         LRMKSLKILWLNGCKRFKSFPEIEVEMESLQVLNMEGIGIRELPRSIANLTGLQSLNLGG
                : :***. * :..*: ::*:**          *::** ****** *   *:**. *.

RMiaGC2         CFNFTR---NPLYCWSALVELDLSENNFVTLPEWISKFVSLKFLHLSDCKSLLEIPQEVL
RMiaGC3         CFNLTRLELRLLHCWSTLQFLDLSGANFVTLPECISKFVSLYVLNLRDCKSLLEIPREVL
RMiaGC1         CFSSPEL--QLLCCWSTLRGLDLSGKNFVTLPECISKFVSLRELDLCDCKSLLEIPQEVL
                **. .    . * ***:*  ****  ******* *******  * * *********.***
```

```
RMiaGC2          PPRVDRVELCNCTSLKKFPKLPLSSGVNFNSLILTNCFRLRGYNITENIVLEQVSSHPHS
RMiaGC3          PPRVGLVFLDNCTSLKNIPELPLSLEVEFKHLSLINCVRLRGYDITENSILDQVSSLPHS
RMiaGC1          PPRILSVCLDNCTSLEKIPKLPPSSEVENRRLSLINCERLRGYDITENSILDQVSSHPHS
                 ***:  * * *****:::*:** *  *: . * * ** *****:**** :*:**** ***

RMiaGC2          QFEITLPGDEVPKWFSCCKDATLVKDEYSSDEEEE------------DEDEYSSEVVGRC
RMiaGC3          QFTITLPGDEVPKWFSCCKDATLVKDEYSSDEEEE--------------DEYS----ARC
RMiaGC1          EFEITLPGDEVPKWFSCCKDATLVKDEYSSDEEEEYSSDEEEEYSSDEEEEYSPKVVARC
                 :* ********************************              :***    .**

RMiaGC2          EVCFEIPPNLDWETLRLVICVVIKGR--GNSCAVNKGWAESCVRINGKMVQSISMW----
RMiaGC3          EVCFEIPPNLDWETLRLVICVVNKGNVCGNECTA-----VISVHINGEWVGRISMG----
RMiaGC1          EVCFEIPPNLDWETLKLVMCVVNKGK-SFFRSKV-----ESHVHINRKMVQSIFMEHCVE
                 ***************.**:*** **.     . .        *.** : *  * *

RMiaGC2          --------------TEESHVALKCFPLLNLNLLQGG-EELTRLQ--QGNMCQVIFPFCEV
RMiaGC3          --------------IEDSNVGLSCIPLLDLRNTAIRWEKLKQLQYVGGNKCQIIFEFSLA
RMiaGC1          FIDRRNELLPSRVLAEETHVALNCIPLLNMSNLRVG-EELTRLQ--QGNMCQIIFEFSGM
                                *:::*.*.*:***::       *:*..**   ** **:** *.

RMiaGC2          PLATPVKILCGVHLLGHQVADVTVDRGQRQWLLLDDDDDDDDDDDDDDDDDDDDDDDQHQ
RMiaGC3          A--TPVKILWGVHLLGHQL-----------------------------------------
RMiaGC1          P--TPVNILCGVHLLGHQVADVTVDRGQRQWLL-----------HDAMAVDDDIHDDQHQ
                 .  ***:** ********:

RMiaGC2          DNELPCLPSASETSLGKRTRSLDFMALDDDHGANIVDVGNHEAQRGEADHPKRRHTDLNE
RMiaGC3          ------LMSASETGLGKGHRLS--------------DVGDHEAQRGEADRPKRRHTDLNE
RMiaGC1          DNELLSLPSASETSLGKRPRLSDFMALDDDHRANVVDVGDHEAQRGEADHPKRRHTDLNE
                       * *****.***  *                ***:*********.**********

RMiaGC2          EPRQ
RMiaGC3          EPKQ
RMiaGC1          EPKQ
                 **.*
```

**Figure S3:** Alignment of the predicted proteins of the three RMia gene candidates. The TNL canonical domains are shown in yellow (TIR), magenta (NB-ARC), red (NLL), blue (LRR) and green (PL-CJiD). Conserved motifs and polymorphic regions in the PL-CJiD domain are highlighted in red and dotted lines, respectively.
